# Fast and Accurate Peptide–MHC Structure Prediction via an Equivariant Diffusion Model

**DOI:** 10.1101/2025.04.28.650973

**Authors:** David Frühbuß, Coos Baakman, Siem Teusink, Erik Bekkers, Stefanie Jegelka, Li C. Xue

## Abstract

Accurate modeling of peptide–MHC (Major Histocompatibility Complex) structures is critical for the development of personalized cancer vaccines and T-cell therapies, as MHC proteins present peptides on the cell surface for immune recognition. Here, we introduce MHC-Diff, a specialized SE(3)-equivariant diffusion model leveraging Geometric Deep Learning to predict the 3D C-alpha atom structures of peptide–MHC complexes with high accuracy. Unlike previous deep learning models that generate a single static structure, our probabilistic approach samples multiple diverse candidates, capturing the inherent flexibility of peptide–MHC binding. Validated on the Pandora benchmark and experimental X-ray crystallography data, MHC-Diff achieves sub-angstrom accuracy, outperforming existing methods by a large margin while matching the inference speed of the fastest available techniques. By enabling rapid and highly accurate structure prediction across diverse peptide lengths and MHC alleles, MHC-Diff provides a powerful new tool for accelerating the design of next-generation cancer vaccines and T-cell therapies.

## 1 Introduction

### Cancer Immunotherapy and the Need for Structural Modeling

Cancer remains a leading cause of mortality world-wide, characterized by high genetic heterogeneity, rapid mutation rates, and the ability to evade immune surveillance. While traditional therapies—surgery, radiation, and chemotherapy—remain standard treatments, they often entail significant side effects and variable outcomes, particularly in late-stage cancers. In contrast, cancer immunotherapy has emerged as a transformative strategy, leveraging the immune system’s innate capacity to recognize and eliminate tumor cells. Techniques such as immune checkpoint blockade and chimeric antigen receptor (CAR) T-cell therapy have demonstrated durable clinical responses, underscoring the potential of immune-based interventions (Pardoll 2012, June et al. 2018).

Central to these therapies is the interaction between T-cell receptors (TCRs) and peptides bound to Major Histocompatibility Complex (MHC) molecules. MHC proteins present intracellular peptides on the cell surface, enabling TCRs to distinguish healthy from abnormal cells. The predictive identification of immunogenic peptides—particularly tumor-specific neoantigens—has therefore become a key step in the design of personalized immunotherapies (Blass and Ott 2021).

However, this predictive task is far from trivial. Human leukocyte antigen (HLA) genes, which encode MHC proteins, are among the most polymorphic loci in the human genome, with over 40,000 known alleles (Robinson et al. 2020). Each allele exhibits unique binding preferences for peptide length, sequence, and conformation. Combined with the combinatorially vast peptide space derived from tumor mutations, this makes the peptide-MHC (pMHC) binding prediction problem a high-dimensional, data-scarce challenge.

### Existing Approaches and Their Limitations

Several computational strategies have been developed to address this challenge. Mass spectrometry (MS)-based approaches can directly identify pMHC pairs in biological samples but are constrained by cost, throughput, and sensitivity (Bassani-Sternberg and Gfeller 2016). In contrast, sequence-based deep learning methods such as NetMHCpan (Jurtz et al. 2017) and MHCflurry (O’Donnell et al. 2020) infer binding affinities from peptide and allele sequences. These models are efficient and widely adopted in clinical pipelines, but often struggle with generalization across rare alleles, peptide length variability, and limited interpretability due to the lack of structural context (Marzella et al. 2024, Atkins et al. 2024).

Recent advances in protein structure prediction, led by AlphaFold2 (Jumper et al. 2021), have inspired structure-based approaches for pMHC modeling. Structural modeling captures spatial and chemical interactions that sequence-based models overlook, offering deeper insights into T-cell recognition and immunogenicity. However, applying models like AlphaFold2 directly to pMHC complexes remains computationally expensive and not optimized for the specific characteristics of peptide-MHC binding. Moreover, these methods typically generate a single structure per input, limiting their ability to represent the conformational flexibility inherent to peptide binding.

### The Promise of Generative and Equivariant Models

Generative models, particularly diffusion models, have shown great promise in molecular and protein structure generation (Trippe et al. 2022, Jing et al. 2023). These models learn the conditional distribution over molecular conformations and can sample diverse, physically plausible structures. Equivariant diffusion models further enhance this capability by respecting geometric symmetries—ensuring that predictions are invariant to rotations and translations—making them particularly well-suited for tasks involving spatial protein structures (Hoogeboom et al. 2022, Schneuing et al. 2022, Zhang et al. 2023). Despite these advances, prior work has not yet delivered a structure prediction model that is both fast and accurate enough for high-throughput personalized immunotherapy. Such a model must (1) generalize across diverse MHC alleles and peptide lengths, (2) match or exceed the accuracy of current structure-based methods, and (3) produce results orders of magnitude faster than AlphaFold-based pipelines to enable population-scale screening.

### Our Approach: MHC-Diff

To address these challenges, we introduce **MHC-Diff**, a conditional SE(3)-equivariant diffusion model for peptide-MHC structure prediction. MHC-Diff takes as input a known MHC pocket structure and a candidate peptide sequence, and generates a distribution of possible C-alpha peptide conformations conditioned on the MHC scaffold. While our model currently predicts C-alpha atom positions, full backbone and sidechain reconstructions can be subsequently achieved with high fidelity (Kryś et al. 2024), given fixed C-alpha geometries.

The model is trained on a curated combination of experimental X-ray structures and physically simulated pMHC complexes, allowing it to learn meaningful geometric and biochemical features of binding. Through design choices such as chain-aware positional encodings and scalable sampling schedules, MHC-Diff achieves competitive accuracy while dramatically reducing inference time.

Compared to previous methods, MHC-Diff offers several practical advantages:

- It supports diverse peptide lengths and MHC alleles, generalizing well to unseen classes.
- It generates multiple candidate structures per input, capturing the flexibility of peptide binding and improving the odds of recovering the native-like conformation.
- It is fast: our model generates 10 diverse candidate structures in under 6 seconds, enabling practical high-throughput applications.
- It maintains high fidelity to experimental X-ray structures, achieving sub-angstrom accuracy on held-out benchmarks.

### Contributions

In this paper, we make the following contributions:

- We develop **MHC-Diff**, a conditional SE(3)-equivariant diffusion model tailored for peptide-MHC structure prediction, capable of generating multiple valid conformations per input. Chain-aware positional encodings and efficient sampling schedules are critical components.
- We evaluate our model on two carefully clustered datasets: an 8K single-allele 9-mer benchmark and a 100K multiallele benchmark spanning multiple peptide lengths, demonstrating strong performance across generalization, accuracy, speed, and robustness.
- We introduce architectural and inference-time optimizations that enable steering the trade-off between generation speed and prediction accuracy.
- We show that our model leverages both experimental and simulation-based data to achieve strong generalization to X-ray targets, even on alleles and peptide lengths not seen during training.
- We release our codebase and curated datasets publicly, facilitating further research in structure-based immunotherapy design.

By bridging the gap between structural accuracy and computational efficiency, MHC-Diff lays the groundwork for nextgeneration immunotherapy pipelines capable of rapid and personalized prediction at scale.

## 2 Method

### 2.1 Diffusion Models

The goal of our model is to predict peptide-MHC structures in their most stable and possibly bound conformation. The structure of the MHC proteins, and specifically the pocket region where the peptides bind (called the G-domain), is fixed. The peptide, on the other hand, is flexible and can adopt multiple possible conformations in the binding pocket. We therefore model each peptide as a probability distribution conditioned on its amino acid sequence and the folded structure and amino acid sequence of the MHC protein’s G-domain. Ultimately, the task of our model is to generate correct joint structures for all peptides and MHCs.

Generative models are the leading paradigm for modeling data distributions and generating new samples from them. Within generative modeling, diffusion models have shown great promise in modeling complex distributions of molecules, which is why we will use them for this problem.

The fundamental idea behind diffusion models and other similar generative models, such as continuous normalizing flows or flow matching, is to learn a transformation from a distribution that we can easily sample from, *p*(*Z*_*T*_), to the true data distribution, *p*(*X*). In diffusion models, we choose this simple distribution to be an isotropic Gaussian, so that we can define this transformation as a stepwise addition of Gaussian noise to the samples from our data distribution until we reach the isotropic Gaussian.

The training objective of our neural network is to learn the inverse of this transition by reversing the noising process, such that we minimize the variational lower bound of the log-likelihood of our data distribution.

This approach allows us, during sampling, to randomly sample from our starting distribution and then stepwise generate joint structures using the posterior of the transitions.

### 2.2 Equivariance

While some recent work has shown that, for large datasets, only a limited amount of geometric inductive bias is required, for most molecule generation problems it is generally beneficial to encode equivariances and invariances into the modeling problem. Equivariance refers to the property of a function whereby, if the input is transformed in some way, the output is transformed correspondingly. Invariance, on the other hand, means that the function’s output remains unchanged under a transformation of the input.

In our neural network, we use SE(3) equivariance to ensure that we learn an invariant target distribution with respect to the group SE(3), which includes rotations and translations in Euclidean space but excludes reflections. For our problem, this is beneficial as the joint molecular structures are inherently three-dimensional structures, and their configurations are invariant under rotations and translations. Furthermore, it has been shown that this approach not only enhances the model’s performance but also reduces the amount of data needed for training.

In our diffusion model, we will use an E(3) equivariant neural network to represent the conditional transition probability *p*(*z*_*t−*1_ |*z*_*t*_) and an isotropic Gaussian, which is inherently O(3) invariant, as the starting distribution *p*(*Z*_*T*_). Given these two conditions, we can prove that the marginal target distribution *p*(*X*) will also be O(3) equivariant.

#### Translational Invariance

We have shown that our model can learn an O(3) invariant distribution using an O(3) invariant starting distribution and an O(3) equivariant conditional transition distribution. However, this does not yet include translation equivariance, even though our neural network is equivariant to both translation and rotation. It is actually not possible to learn a translation-invariant distribution in ℝ^3^. The reason for this is that such a distribution would have to satisfy *p*(*x* + *t*) = *p*(*x*) for any translation *t* in ℝ^3^, which means it would have to be constant. If it were constant, it could no longer integrate to 1 and therefore could not be a probability distribution.

Koller et al. (Köhler et al. 2020) was the first to propose limiting the modeling problem to a subspace that satisfies ∑*x* =| 0. Technically, this constraint means that positions lie on an (*M −* 1)-dimensional subspace with *x* ∈ ℝ^*M ×N*^, but in practice, we simply use the normal representation space and ensure that all positions and positional noise are centered at the origin, ∑*x* = 0.

#### E(3) Equivariant Graph Neural Network

As mentioned above, we will use an E(3) equivariant neural network to learn the conditional transition probability *p*(*z*_*t−*1_ | *z*_*t*_). There are multiple options to achieve E(3) equivariance, but we will focus on the message-passing framework and the Equivariant Graph Neural Network (EGNN) architecture for our model. It is *E*(3) equivariant because it maintains the structure’s geometric properties throughout the network layers. The design of the message, position, and feature updates ensures that transformations in the input space (such as rotations and translations) lead to predictable and consistent transformations in the output space. These updates ensure that the learned conditional distribution *p*(*z*_*t−*1_ | *z*_*t*_) remains equivariant under *E*(3) transformations.

In combination with the translational invariance and an isotropic Gaussian as the starting distribution *p*(*Z*_*T*_), this will allow our equivariant diffusion model to learn an E(3) invariant target distribution. This provides all the necessary tools to construct our model for generating conditional peptide-MHC structures.

### 2.3 Peptide-MHC structure prediction

Our goal is to predict the joint structure of a peptide, which lacks a known structure, and a specific MHC protein for which the structure is known. To achieve this, we will train a diffusion model with an equivariant graph neural network backbone to generate the structure of the peptide within the protein’s pocket.

#### Input Representation

In this work, we model both the peptide and the protein at their amino acid level. This means we represent the spatial location of each amino acid as a single node. For the protein, we will only use the protein pocket to reduce the number of nodes and thus the computational cost. We represent the location of an amino acid in space as the position of the C-alpha atom of that amino acid. For the protein pocket amino acids, we already know the positions, while for the peptide amino acids, we do not.

In our datasets, we have joint structures of peptides with MHC proteins. Let **z**_data_ = [**x, h**] be a peptide-MHC complex with node positions **x** and node features **h**. Each **x** and **h** consists of nodes from the peptide [**x**_mol_, **h**_mol_] and from the protein pocket [**x**_pro_, **h**_pro_]. Importantly, the peptide features *h*_mol_ will remain fixed in the noising process. For each node *i*, the feature *h*_*i*_ consists of a one-hot encoding of the amino acid type. For the peptide nodes, we additionally add a sinusoidal positional encoding of the amino acid’s position within the chain:

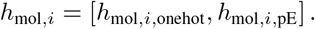

Since the peptide starts as a random point cloud sampled from an isotropic Gaussian distribution, it contains no inherent structural information about the chain connectivity. To guide the model in reconstructing the correct linear structure of the peptide, we embed positional information into the node features using a sinusoidal chain positional encoding (Vaswani et al. 2017). This provides each amino acid node with information about its relative position within the sequence, enabling the model to more easily reconstruct chain-like conformations during denoising. The exact form of the positional encoding is given in Appendix A.

#### Center of Mass Handling

As discussed in Section 2.2, we need our model to be invariant to translations. To achieve this, we restrict the entire modeling problem to a subspace where the peptide is always centered around the origin. Before we start generating noised samples, we subtract the mean of the peptide positions from the entire peptide-MHC complex positions. This moves the peptide to the origin of our coordinate system and adjusts the protein pocket accordingly. Furthermore, we center every noise that we add during training and sampling around the origin and recenter the entire p-MHC complex after every noising and denoising step.

#### The Diffusion Process

The goal of the diffusion process is to train a neural network to perform a sequential stepwise denoising process, starting with a random point cloud and ending with a correctly generated peptide structure.

We can define the noise process as a Markov chain that gradually adds Gaussian noise to the data:

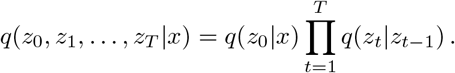

A very important property of this noise process is that it admits sampling every *x*_*t*_ in closed form as:

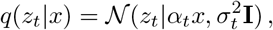

with (*α*_*t*_, *σ*_*t*_). This noise process starts at *x*_0_ (where *α*_0_ = 1 and *σ*_0_ = 0) and ends at the trivial noise distribution 𝒩 (0, 1) (where *α*_*T*_ = 0 and *σ*_*T*_ = 1). The dynamics of the noise process are defined by the noise schedule (*α*_0_, …, *α*_*T*_ and *σ*_0_, …, *σ*_*T*_). There are many important design choices in choosing the dynamics of *α*_*t*_ and *σ*_*t*_, as well as the relative scale of the data and the noise, which are discussed in Section 5.2.

#### Training the Neural Network

The direct sampling property above allows us to train our neural network efficiently on single-step transitions from timestep *t* → 0. This circumvents the error accumulation that would occur when trying to predict the single-step transitions sequentially during training and guarantees equal training of all transition steps. The role of the neural network is therefore to predict *x*_0_ given *x*_*t*_. This can be done by predicting the mean and the variance of *x*_0_ or by predicting the added noise *ϵ*. In practice, predicting *ϵ* is more stable (Ho et al. 2020).

In each forward pass, we therefore sample a timestep *t ∼* U(1, 2, …, *T*) for each peptide-MHC and compute the noised sample:

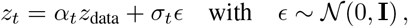

Importantly the node features are not noised in *z*_*t*_. Before we input the noised sample into the neural network, we append the normalized timestep *t/T* to the node features **h**_mol_ and **h**_pro_. We then predict 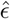 with the neural network. We can try to judge the single-step denoising property by computing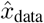:

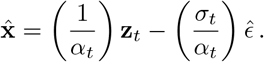

But for training the neural network, we directly use *ϵ*. One interesting note here is that this training is very sparse, considering that for each epoch the model only sees a single timestep for each peptide-MHC.

#### Training Objective

The training objective of the model is to minimize the variational lower bound of the model likelihood:

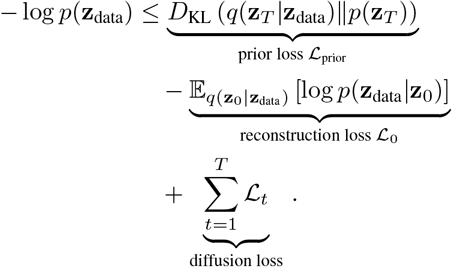

Using the noise prediction parametrization, Kingma et al. (Kingma et al. 2021) has shown that the *L*_*t*_ loss simplifies to:

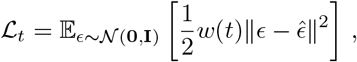

with

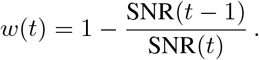

Ho et al. (Ho et al. 2020) has proposed that setting *w*(*t*) = 1 for images during training stabilizes training, which Hoogebloom et al. (Hoogeboom et al. 2022) confirms also holds for molecules. Further, they have shown that for continuous *x, L*_0_ can be computed as:

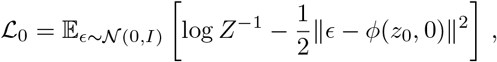

with

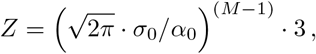

where *M* is the number of peptide nodes.

#### The Generative Denoising Process

During sampling, we are given the full information about the protein pocket [**x**_pro_, **h**_pro_] but only have the feature information **h**_mol_ for the peptide. We record the center of the protein pocket before centering it around the origin. We then initialize the peptide positions **x**_mol_ at the origin.

We use the posterior of the transitions (Kingma et al. 2021) conditioned on the previous timestep to stepwise generate the positions of the peptide nodes in relation to the protein pocket:

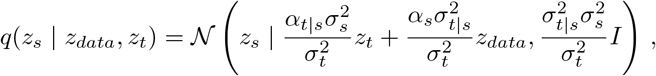

with

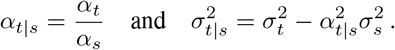

Importantly, after every denoising step, we recenter the peptide to ensure that we do not leave the subspace that our neural network is trained on. Once we reach **z**_0_, we use *q*(*x*_*sampled*_|*z*_0_) to predict the final *x* structure. Then we move the entire peptide-MHC complex back to the original protein pocket center to compare it to our target structure.

### 2.4 Equivariant Neural Network

For our neural network, we use an E(3) equivariant graph neural network, as discussed in Section 2.2. Following Schneuing et al. (Schneuing et al. 2022) we use a modification that makes the model SE(3). This removes equivariance to reflections, which can be problematic for molecules, as two molecules that are reflections of each other can have different properties. The modification is an additive term to the EGNN’s coordinated update:

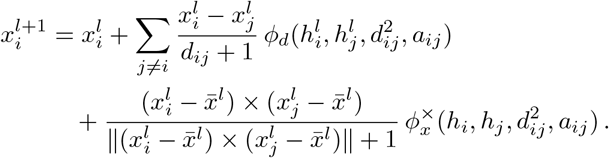

using the fact that the cross product changes sign under reflection. Here, *d*_*ij*_ stands for the pairwise distance between *x*_*i*_ and *x*_*j*_. For graph connectivity, we use a fully connected graph with edge cutoffs. The peptide nodes are fully connected to each other. For protein-protein node interactions, we use a cutoff of 8 Å, and for protein-peptide node interactions, a cutoff of 14 Å. These values are chosen based on experience in modeling peptide-MHC complexes and our observations during training runs. They are intended to incorporate relevant information from nearby nodes while reducing computational cost by removing edges that do not contribute additional interaction information.

The network receives the noised sample *z*_*t*_, which consists of the noised peptide positions, fixed peptide features (including the one-hot amino acid type encoding and the positional encoding), and the normalized time step *t/T*. Additionally, *z*_*t*_ includes the protein pocket [**x**_pro_, **h**_pro_], which is not affected by noise but whose positions adjust with the center of mass centering to maintain its relative position to the peptide. The neural network then predicts the noise for the peptide positions, denoted as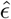.

## 3 Related Work

Several approaches have tackled the challenge of pMHC structure prediction, each advancing the field from different angles.

Early fine-tuning efforts on AlphaFold2 demonstrated that anchor-free peptide placement in the MHC groove was possible (Motmaen et al. 2023). These models achieve high accuracy but retain significant computational overhead and typically predict only a single structure per input.

APE-Gen2.0 introduced anchor-guided sampling and energy minimization to generate ensembles of peptide conformations, including noncanonical geometries and post-translational modifications (Fasoulis et al. 2024). However, APE-Gen remains a physics-based sampler rather than a learned generative model.

PANDORA proposed a template-based approach that threads peptides into MHC pockets based on predicted anchor residues, using loop modeling for flexibility (Marzella et al. 2022). It is computationally efficient but limited by template availability and by evaluating only a single top-scoring pose per input.

SwiftMHC combined an attention-based deep learning framework to simultaneously predict peptide–MHC binding and detailed all-atom structures (Baakman et al. 2025). It achieves high throughput, but does not explicitly model the full conformational distribution.

MHC-Fold integrated geometric deep learning to jointly predict pMHC structures and binding specificity (Aronson et al. 2022), demonstrating that learned representations of molecular geometry can greatly enhance generalization, especially across unseen alleles.

While APE-Gen2.0 and PANDORA generate multiple candidate poses, they do not explicitly learn the distribution of peptide conformations. Our work builds on these foundations by developing a conditional generative model that both accelerates prediction and robustly captures conformational diversity.

## 4 Results

To evaluate the performance and practical utility of **MHC-Diff**, we conducted experiments on two curated datasets designed to assess both predictive accuracy and generalization across MHC alleles and peptide lengths. Our evaluation also emphasizes computational efficiency, a key factor for enabling highthroughput structure prediction in immunotherapy applications.

### 4.1 Datasets and Evaluation Protocol

#### 8K Dataset (Fixed Allele and Peptide Length)

This dataset consists of 7,726 peptide–MHC structures centered on the most common human allele, **HLA-A*02:01**, with all peptides fixed at 9 amino acids in length. It includes 202 experimental X-ray structures from the Protein Data Bank and 7,524 modeled structures from Pandora. To ensure rigorous cross-validation, we clustered the peptides using Gibbs sampling into 10 clusters and performed leave-one-cluster-out cross-validation. This guarantees that peptides in the test set share no close sequence or structural similarity with those in training.

#### 100K Dataset (Diverse Alleles and Peptide Lengths)

To evaluate generalization, we compiled a large-scale dataset spanning diverse MHC alleles and peptide lengths. The dataset includes approximately 100,000 peptide–MHC structures from the Pandora database and 1,000 additional X-ray structures from IMGT. We applied hierarchical clustering over the MHC alleles to form 10 allele-based clusters. One cluster was reserved for validation and two for testing, resulting in a test set of 4,007 samples, including 72 experimentally resolved structures. This setup ensures that all MHC alleles in the test set are entirely unseen during training.

#### Evaluation Metrics

We report four standard metrics based on C_*α*_-RMSD between predicted and experimental peptide conformations:

- **Mean RMSD:** Average across all 10 samples generated per input.
- **Median RMSD:** Typical accuracy of generated structures.
- **Best-of-10 RMSD:** Minimum RMSD across candidates; reflects the model’s capacity to capture near-native states.
- **Highest Outlier RMSD:** Maximum deviation; serves as a robustness indicator.

### 4.2 Performance on Canonical Peptide–MHC Structures (8K Dataset)

Table 1 summarizes the predictive accuracy of all evaluated models on the 8K dataset. MHC-Diff achieves a best-of-10 C*α*-RMSD of 0.45 Å, significantly outperforming all baselines. This metric reflects our model’s ability to generate at least one structure that closely matches the experimentally determined conformation for each peptide–MHC complex.

**Table 1:**
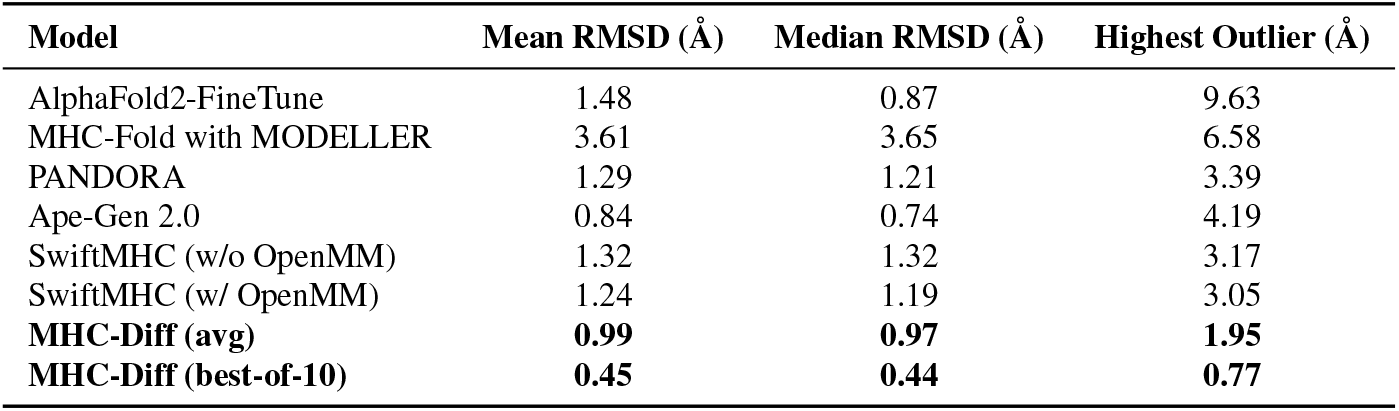
Structure prediction accuracy on the 8K dataset. All methods were evaluated on the same set of 202 X-ray peptide–MHC structures from the HLA-A*02:01 allele with 9-mer peptides. Ensemble-based methods (Pandora, Ape-Gen 2.0) report the lowest RMSD among their respective candidate sets. MHC-Diff generates 10 samples per input; we report both the average RMSD across all candidates and the best-of-10 RMSD to capture both generative precision and robustness.

| Model | Mean RMSD (Å) | Median RMSD (Å) | Highest Outlier (Å) |
| --- | --- | --- | --- |
| AlphaFold2-FineTune | 1.48 | 0.87 | 9.63 |
| MHC-Fold with MODELLER | 3.61 | 3.65 | 6.58 |
| PANDORA | 1.29 | 1.21 | 3.39 |
| Ape-Gen 2.0 | 0.84 | 0.74 | 4.19 |
| SwiftMHC (w/o OpenMM) | 1.32 | 1.32 | 3.17 |
| SwiftMHC (w/ OpenMM) | 1.24 | 1.19 | 3.05 |
| MHC-Diff (avg) | **0.99** | **0.97** | **1.95** |
| MHC-Diff (best-of-10) | **0.45** | **0.44** | **0.77** |

Although multiple conformations can be valid for a given complex, only one X-ray structure is available per test instance. Therefore, the best-of-10 metric provides a practical upper bound on model precision—assessing whether our generative approach can sample a near-native structure. To complement this, we also report the average RMSD across all generated candidates, which reflects overall prediction quality and structural diversity. MHC-Diff maintains competitive performance even in this more challenging setting, with an average RMSD of 0.99 Å.

Deterministic models such as AlphaFold2-FineTune and MHC-Fold predict only a single structure per input. In contrast, methods like Pandora and Ape-Gen 2.0 generate fixed-size ensembles of candidate conformations. For these, we report the lowest RMSD within the provided ensemble to enable fair comparison to MHC-Diff’s generative predictions. However, these methods lack adaptive diversity and do not explicitly model the distribution over valid conformations.

We also note that AlphaFold2-FineTune and MHC-Fold were not retrained on our curated 8K dataset due to computational constraints. As a result, some data leakage from the test set into the pretraining corpus cannot be ruled out, which may inflate their apparent accuracy.

As shown in Figure 3, MHC-Diff not only delivers the lowest prediction errors on average, but also exhibits greater robustness across individual test structures. While other methods produce extreme outliers—such as AlphaFold2-FineTune with a 9.60 Å maximum error—MHC-Diff’s worst-case prediction remains below 2 Å.

**Figure 1:**
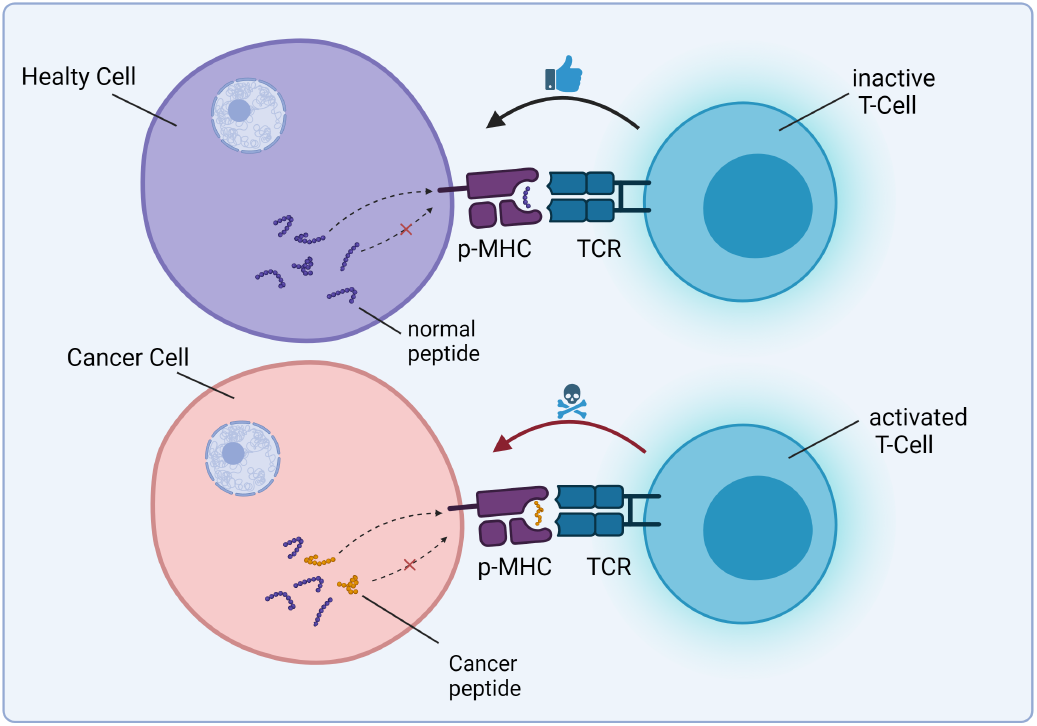
Peptide–MHC Presentation and T-Cell Activation. Intra-cellular peptides, including cancer-specific neoantigens, are generated during cellular metabolism and presented on the cell surface by Major Histocompatibility Complex (MHC) proteins. T-cells continuously scan these peptide–MHC (pMHC) complexes via their T-cell receptors (TCRs), enabling the detection of abnormal cells. The structural properties of pMHC complexes are critical for immune recognition and response, underscoring the importance of accurate pMHC structure prediction for cancer immunotherapy development.

**Figure 2:**
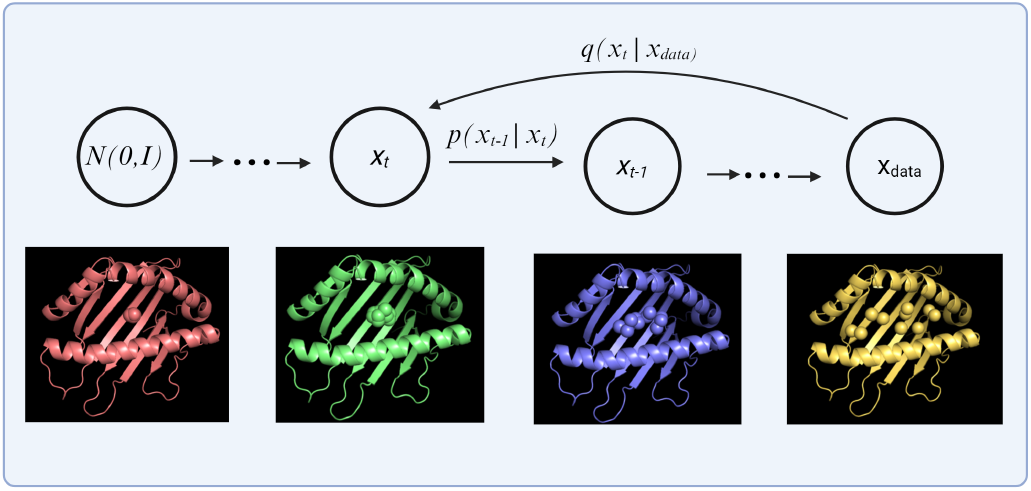
Overview of Diffusion for peptide-MHC: During training we noise our structures to train our neural network to predict which part of the structure is noise. During sampling we use our trained neural network to denoise from random noise to our target peptide structure.

**Figure 3:**
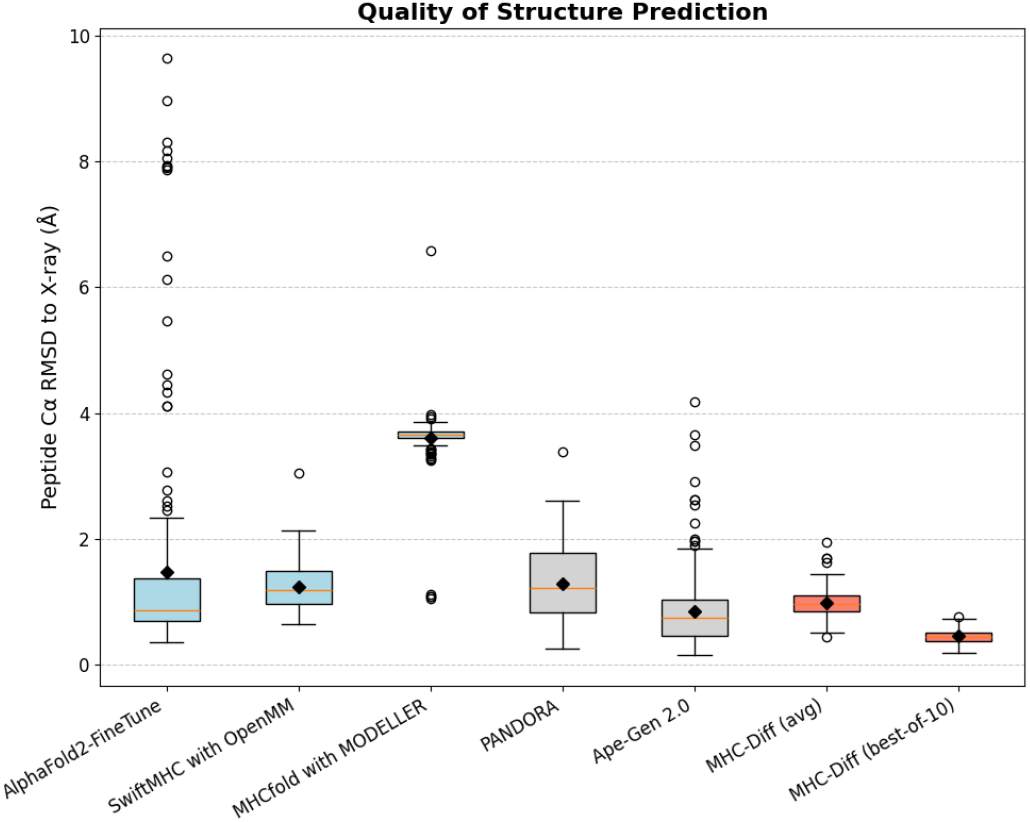
Prediction accuracy on 202 X-ray peptide–MHC complexes from the 8K dataset. We report the C*α*-RMSD for each individual structure in the test set. These test structures were held out using a leave-one-cluster-out cross-validation scheme, where clustering was performed based on peptide similarity across the full 8K dataset (HLA-A*02:01, 9-mer peptides). This ensures that test structures are dissimilar in sequence from any training example. MHC-Diff achieves consistently lower RMSD values and reduced variance, with significantly fewer high-error predictions compared to other methods. Each circle represents an individual experimentally resolved peptide–MHC complex that was identified as a high-error outlier.

### 4.3 Generalization Across Diverse Alleles and Peptide Lengths (100K Dataset)

We next evaluate MHC-Diff on the 100K dataset, which spans a broad range of MHC alleles and peptide lengths. Performance is assessed on 72 experimental X-ray complexes drawn from the held-out clusters of unseen alleles, providing a goldstandard evaluation of generalization.

Despite the increased structural diversity relative to the 8K benchmark, MHC-Diff achieves a best-of-10 C_*α*_-RMSD of 0.61 Å and a mean C_*α*_-RMSD of 1.50 Å (Table 2). An overlay of a predicted structure against its corresponding X-ray target is shown in Figure 6, illustrating the model’s capacity to recover fine-grained backbone conformations even for unseen alleles and longer peptides.

**Table 2:** Generalization performance of MHC-Diff on the 100K dataset, evaluated on 72 unseen experimental X-ray structures spanning diverse MHC alleles and peptide lengths.

| Model | Mean RMSD (Å) | Best-of-10 RMSD (Å) |
| --- | --- | --- |
| MHC-Diff (ours) | 1.50 | 0.61 |

To further analyze the effect of peptide length, we report results stratified by peptide size (Tables 3 and 4). For 8-mer peptides—the most common length in the dataset—MHC-Diff achieves a best-of-10 RMSD of 0.53 Å, with longer peptides (9–10 residues) showing a slight increase in error, as expected due to greater conformational variability. Even for peptides of length 11–13, where only a single test structure is available for each length, the model maintains sub-angstrom best-of-10 accuracy.

**Table 3:** Best-of-10 RMSD grouped by peptide length for MHC-Diff predictions on the 100K dataset (72 X-ray structures).

| Peptide Length | N (X-ray Structures) | Best-of-10 RMSD (Å) | Range (Å) |
| --- | --- | --- | --- |
| 8 | 37 | 0.53 | 0.34–0.72 |
| 9 | 23 | 0.70 | 0.28–1.44 |
| 10 | 9 | 0.74 | 0.45–1.46 |
| 11 | 1 | 0.48 | – |
| 12 | 1 | 0.36 | – |
| 13 | 1 | 0.58 | – |

**Table 4:** Mean RMSD by peptide length, quantifying ensemble diversity on the 100K dataset (72 X-ray structures).

| Peptide Length | N (X-ray Structures) | Mean RMSD (Å) | Range (Å) |
| --- | --- | --- | --- |
| 8 | 37 | 1.40 | 0.85–2.48 |
| 9 | 23 | 1.60 | 1.09–2.42 |
| 10 | 9 | 1.48 | 1.10–1.89 |
| 11 | 1 | 1.78 | – |
| 12 | 1 | 2.34 | – |
| 13 | 1 | 2.54 | – |

Overall, these results demonstrate that MHC-Diff generalizes robustly across peptide lengths and allele diversity, maintaining sub-angstrom candidate recovery in most cases. Notably, the best-of-10 RMSD on the 100K dataset (0.61 Å) remains lower than the best performance of competing methods on the much simpler 8K benchmark (e.g., Ape-Gen 2.0 at 0.84 Å), highlighting the substantial improvement in both accuracy and generalization enabled by our approach.

### 4.4 Speed and Efficiency

Rapid structure generation is critical for large-scale screening of peptide-MHC pairs. Figure 4 compares generation times across different methods. For MHC-Diff, the reported time of 5.6 seconds corresponds to generating 10 candidate conformations per input, or approximately 0.56 seconds per structure. All timings include structure saving to disk and were measured at each model’s optimal batch size (e.g., SwiftMHC at 60, MHC-Diff at 100) to ensure consistent comparison. A detailed breakdown of times can be found in Appendix 5.

**Figure 4:**
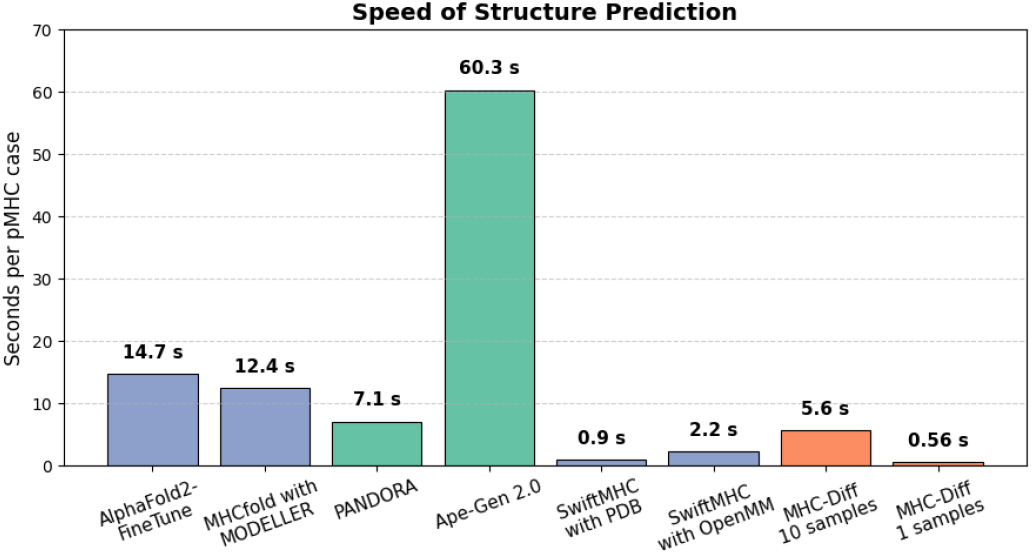
Generation speed across methods. Sampling times are measured at optimal batch sizes and include time for saving output structures. Full numeric results are available in the appendix.

It is important to note that MHC-Diff generates only C*α*-atom coordinates. Full backbone and sidechain reconstruction, while not performed natively, can be efficiently achieved using external tools such as PDBfixer or MODELLER. Based on typical tool performance, this postprocessing would add approximately 1 second per structure. Even when accounting for this step, MHC-Diff remains among the fastest available methods for peptide-MHC structure prediction.

In addition to its high base speed, MHC-Diff offers flexible control over generation time by adjusting sampling parameters. Figure 5 shows how reducing the number of diffusion sampling steps accelerates generation with modest accuracy tradeoffs. For instance, reducing sampling steps from 1000 to 200 lowers generation time from 5.8 to 2.1 seconds for 10 conformations, while still maintaining sub-angstrom best-of-10 RMSD. Reducing further to 125 steps increases speed to 1.7 seconds but results in a larger drop in average RMSD. Full tabulated results are provided in Appendix 6.

**Figure 5:**
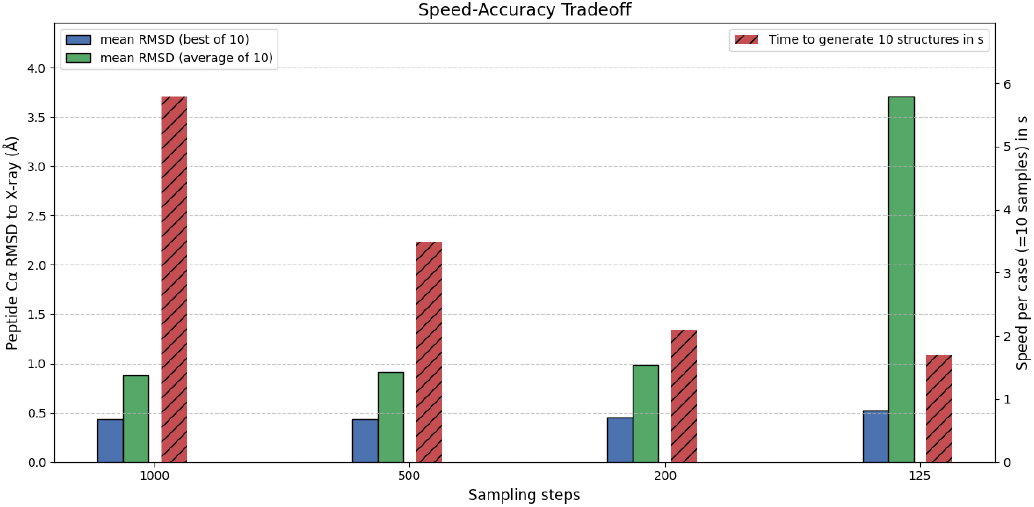
Sampling speed-accuracy tradeoff for MHC-Diff. Reducing the number of sampling steps decreases generation time while maintaining sub-angstrom best-of-10 RMSD up to 200 steps. Full numeric results are provided in the appendix.

**Figure 6:**
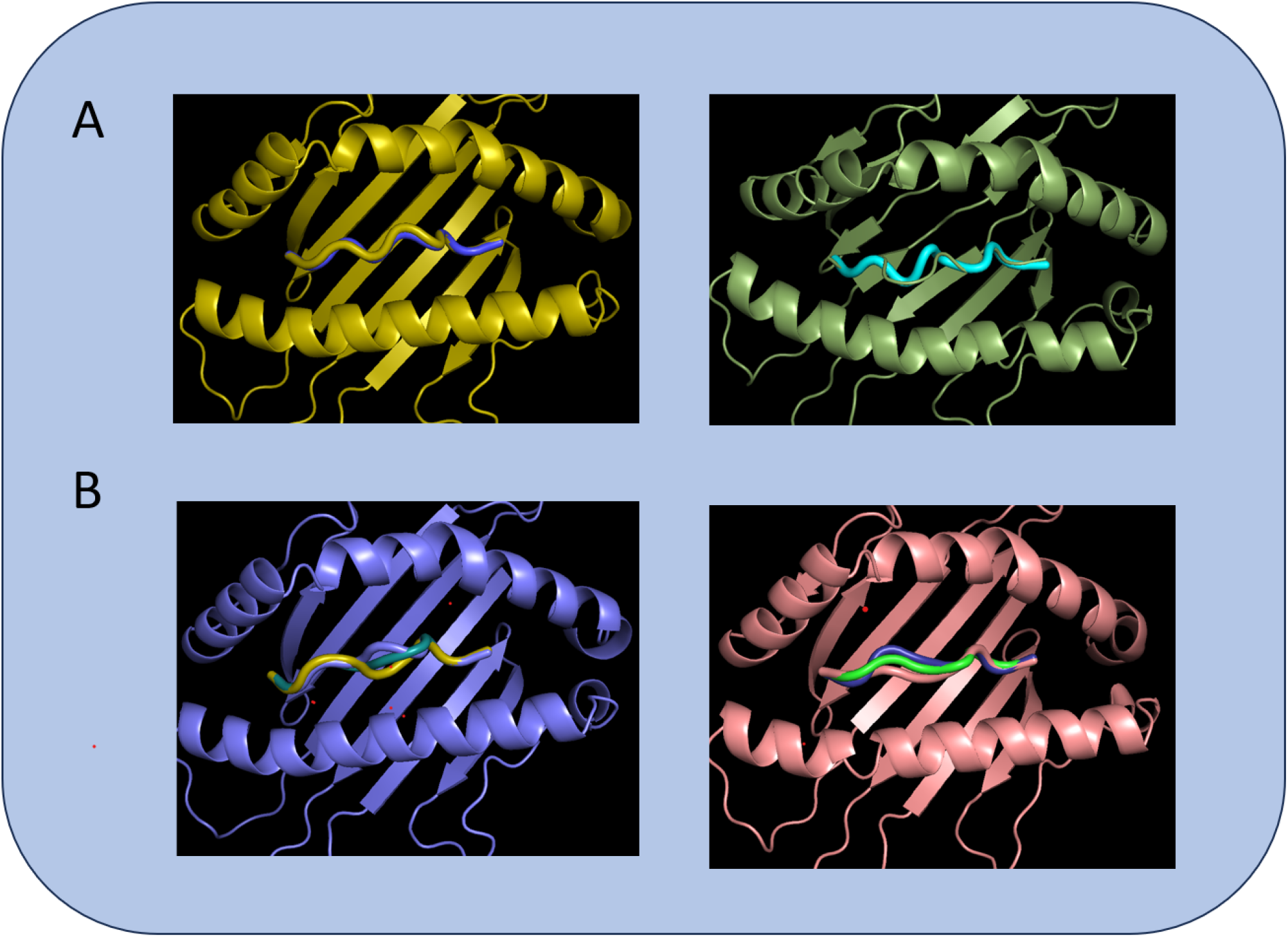
Qualitative evaluation of predicted peptide–MHC structures. (A) Comparison of predicted structures to experimental targets. Left: Structure of a 9-mer peptide bound to HLA-A*02:01 (PDB ID: 3V5D) from the 8K dataset, showing the experimental structure (olive) and our prediction (blue). Right: Structure of a 10-mer peptide bound to HLA-A*02:01 (PDB ID: 3DX7) from the 100K dataset, with the experimental structure (green) and our prediction (turquoise). (B) Structural diversity captured by the model. Left: Multiple predicted conformations for the 9-mer peptide from PDB ID: 1AKJ (8K dataset). Right: Multiple predicted conformations for the 10-mer peptide from PDB ID: 3PAB (100K dataset). The predicted structures demonstrate both high accuracy relative to crystallographic targets and the ability to model realistic conformational flexibility.

Even when accounting for backbone and sidechain recovery, MHC-Diff enables rapid peptide-MHC structure generation, achieving sub-2 second runtimes per full-atom model if desired. This high speed makes it practical for large-scale applications such as binding affinity prediction or immunogenicity screening, where millions of structures may need to be evaluated.

### 4.5 Generative Diversity and Structural Flexibility

Accurately modeling the structural flexibility of peptide-MHC (pMHC) complexes is critical for downstream applications such as T-cell receptor (TCR) recognition and binding affinity prediction. Unlike methods that predict a single static structure, MHC-Diff explicitly learns a distribution over plausible peptide conformations conditioned on the MHC context.

While experimental datasets typically provide only one conformation per peptide-MHC complex, biological evidence suggests that multiple binding modes are possible. As a result, direct evaluation of generative diversity is challenging. Nevertheless, MHC-Diff achieves low average RMSD values across generated samples—0.88 Å on the 8K benchmark and 1.50 Å on the 100K dataset—indicating that most candidate structures are close to the experimental ground truth even without selecting the best one.

Furthermore, expert visual inspection shows that generated structures consistently maintain correct anchoring within the MHC binding groove, even when RMSD values are moderately higher. This qualitative agreement provides additional evidence that MHC-Diff learns a biologically meaningful distribution rather than overfitting to a single conformation.

Quantitatively, over 88% of examples include at least one generated structure that closely matches the X-ray conformation, while approximately 12% of candidates diverge significantly (e.g., the peptide placed outside the MHC pocket). Such outliers are easily removed using simple RMSD or pocket occupancy filters.

## 5 Discussion

The ability to accurately and rapidly model peptide-MHC (pMHC) complexes is a critical bottleneck in immunotherapy development. MHC-Diff addresses this challenge by offering a fast, high-fidelity, and generalizable structure generation framework. Compared to previous structure-based methods, MHC-Diff achieves sub-angstrom accuracy on held-out targets while maintaining high generalization across diverse MHC alleles and peptide lengths. In terms of throughput, only SwiftMHC and MHCFold approach similar speeds, but unlike these methods, MHC-Diff explicitly samples diverse peptide conformations.

Capturing conformational diversity is particularly important for improving predictions of TCR engagement (Borbulevych et al. 2009, Hawse et al. 2013), a major determinant of successful T-cell–based therapies and neoantigen vaccine efficacy (Abd-Aziz and Poh 2022, Liu et al. 2021). By modeling the range of plausible peptide structures within the MHC binding groove, MHC-Diff lays the groundwork for better understanding and engineering of immune responses in cancer immunotherapy. While recent large models such as AlphaFold 3 (Abramson et al. 2024) and Boltz 1 (Wohlwend et al. 2024) can also capture conformational diversity, we do not benchmark against them here, as they are orders of magnitude slower than MHC-Diff without clear accuracy gains for pMHC modeling based on their own claims.

### 5.1 Limitations

While MHC-Diff significantly advances pMHC modeling, some limitations remain. Currently, the model predicts only C-alpha atom positions, leaving full backbone and sidechain reconstruction to downstream tools, allowing users to flexibly balance speed and accuracy.

Our method also benefits from pre-aligning structures to a reference frame during training, a minimal preprocessing step specific to our positional encoding design. Finally, rare sampling instabilities can occur if trajectories drift from the learned manifold, as discussed in the Appendix.

### 5.2 Future Work

Looking forward, an important direction is incorporating MHC-Diff into a full pMHC binding prediction framework, such as DeepRank (Crocioni et al. 2024). Additional improvements could involve targeted energy minimization for studying TCR-pMHC interactions, extending the method to MHC class II complexes, and applying the same design principles to other critical structure generation tasks requiring both high accuracy and speed.

By bridging structural accuracy, conformational diversity, and computational efficiency, MHC-Diff lays the groundwork for the next generation of structure-driven immunotherapy pipelines.

## Appendix Overview

This appendix provides additional experimental details, full technical descriptions, and proofs supporting the methods introduced in the main text. It is organized as follows:

- Additional tables with full breakdowns of model performance and generation speed.
- Dataset preprocessing protocols and allele/length distributions.
- Mathematical foundations: types of equivariances, proofs of model invariances, and SE(3)-equivariant neural network architecture.
- Input representation choices, noise schedule designs, and detailed handling of sampling and positional encoding. Where appropriate, figures and tables are referenced directly in the main text.

## A Supplementary Results

Table 3 reports structure prediction accuracy (Best-of-10 RMSD) stratified by peptide length on the 100K dataset.

Table 4 shows the ensemble diversity measured as Mean RMSD across the generated candidates.

Table 5 compares generation speed between MHC-Diff and baseline methods.

**Table 5:** Generation speed comparison. MHC-Diff offers competitive throughput while maintaining high structural accuracy.

| Method | Time per Structure (s) |
| --- | --- |
| AlphaFold2-FineTune (with MODELLER) | 14.70 |
| MHCfold | 12.40 |
| PANDORA | 7.10 |
| Ape-Gen 2.0 | 60.30 |
| SwiftMHC (with PDB) | 0.90 |
| SwiftMHC (with OpenMM) | 2.20 |
| **MHC-Diff (10 candidates)** | 5.60 |
| **MHC-Diff (1 candidate)** | 0.56 |

Table 6 quantifies the speed-accuracy tradeoff achievable by adjusting the number of diffusion steps.

**Table 6:** Speed-accuracy tradeoff for MHC-Diff. Fewer steps yield faster generation at the cost of minor accuracy degradation.

| Sampling Steps | Best RMSD (Å) | Mean RMSD (Å) | Time (s) |
| --- | --- | --- | --- |
| 1000 | 0.44 | 0.88 | 5.80 |
| 500 | 0.43 | 0.91 | 3.50 |
| 200 | 0.45 | 0.99 | 2.10 |
| 125 | 0.52 | 3.71 | 1.70 |

## B Dataset Preprocessing and Overview

To enable robust training and benchmarking, we curate two peptide–MHC datasets combining experimental X-ray complexes (Lefranc et al. 2005, Berman et al. 2000) and modeled structures (Marzella et al. 2022). Specific preprocessing steps include:

- **Structure Clustering:** Different clustering strategies were applied depending on the dataset. For the **8K dataset** (fixed HLA-A*02:01 allele and fixed peptide length), peptides were clustered based on sequence similarity using Gibbs sampling, ensuring no closely related peptide sequences are shared between training and test folds. For the **100K dataset** (diverse alleles and peptide lengths), hierarchical clustering was performed over the MHC alleles based on pocket similarity. Test and validation splits were constructed by holding out entire allele clusters, ensuring that alleles in the test set were entirely unseen during training. The alleles held out for testing were: MH1-K1b, HLA-C*0501, HLA-C*0602, H2-D1*02, RT1-A, HLA-B*4403, HLA-B*4402, MH1*0401, MH1-N*01301, HLA-A*0205, HLA-B*3508, HLA-B*0801, and MH1-A*01.
- **Alignment:** All peptide–MHC structures were rigidly aligned to a reference MHC pocket frame to standardize coordinate systems across the dataset. This stabilization improves both model convergence and the effectiveness of positional encoding.
- **Pocket Extraction:** Only residues forming the MHC binding groove were retained. Removing the rest of the MHC structure reduces computational complexity and focuses learning on the biologically relevant peptide-binding site.
- **Peptide Representation:** Peptides were modeled at the residue level, with each amino acid represented by its C-alpha atom position. This coarse-grained representation balances predictive performance with computational efficiency.

## C Types of Equivariances

There are various types of equivariances relevant to molecular modeling:

### Orthogonal Group

*O*(3) The orthogonal group *O*(3) consists of all 3×3 orthogonal matrices. These matrices represent rotations and reflections in three-dimensional space. A function *f* is said to be equivariant with respect to *O*(3) if:

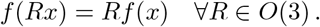

where *x* is the input vector and *R* is an orthogonal transformation. Equivariance to *O*(3) ensures that the function handles rotations and reflections consistently.

### Special Orthogonal Group

*SO*(3) The special orthogonal group *SO*(3) is a subgroup of *O*(3) consisting of all 3×3 orthogonal matrices with determinant 1. These matrices represent rotations in three-dimensional space, but not reflections. A function *f* is said to be equivariant with respect to *SO*(3) if:

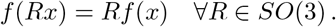

Equivariance to *SO*(3) is particularly important for models that need to account for rotations but not reflections, such as those used in protein structure prediction.

### Euclidean Group

*E*(3) The Euclidean group *E*(3) extends *SO*(3) by including translations in addition to rotations. An element of *E*(3) is represented as a pair (*R, t*), where *R* ∈ *SO*(3) and *t* ∈ ℝ^3^. A function *f* is equivariant with respect to *E*(3) if:

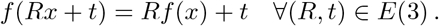

Equivariance to *E*(3) is crucial for models dealing with spatial transformations that include both rotations and translations, such as those used in the prediction of molecular structures in three-dimensional space.

### Special Euclidean Group

*SE*(3) The special Euclidean group *SE*(3) is a specific instance of *E*(3) focused on rigid body transformations. It is represented as a pair (*R, t*), where *R* ∈ *SO*(3) and *t* ∈ ℝ^3^. A function *f* is equivariant with respect to *SE*(3) if:

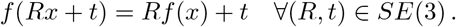

## D Invariant Marginal Distribution

In our diffusion model, we will use an *E*(3) equivariant neural network to represent the conditional transition probability *p*(*z*_*t−*1_ | *z*_*t*_) and an isotropic Gaussian, which is inherently *O*(3) invariant, as the starting distribution *p*(*Z*_*T*_). Given these two conditions, we can prove that the marginal target distribution *p*(*X*) will also be *O*(3) equivariant (Xu et al. 2022).

### Proof of Invariant Marginal Distribution

- Base case: Observe that *p*(*z*_*T*_) = *N* (0, *I*) is equivariant with respect to rotations and reflections, so *p*(*z*_*T*_) = *p*(*Rz*_*T*_).
- Induction step: For some *t* ∈ {1, …, *T*}, assume *p*(*z*_*t*_) to be invariant meaning that *p*(*z*_*t*_) = *p*(*Rz*_*t*_) for all orthogonal *R*. Let *p*(*z*_*t−*1_|*z*_*t*_) be equivariant meaning that *p*(*z*_*t−*1_|*z*_*t*_) = *p*(*Rz*_*t−*1_|*Rz*_*t*_) for orthogonal *R*. Then:

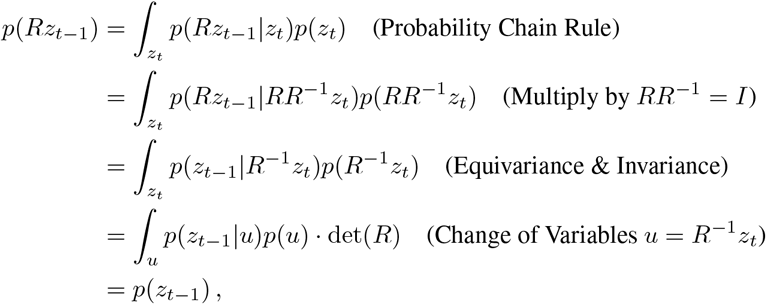

and thus *p*(*z*_*t−*1_) is invariant. By induction, *p*(*z*_*T −*1_), …, *p*(*z*_0_) are all invariant.

## E Equivariant Graph Neural network

The EGNN architecture achieves E(3) equivariance through its update rules:

### Message Computation

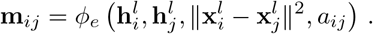

Messages **m**_*ij*_ are generated using node features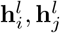, the squared Euclidean distance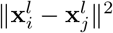, and edge features *a*_*ij*_. The use of the squared distance, which is invariant under rotations and translations, ensures that the computed messages are invariant to these transformations.

### Position Update

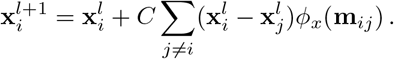

Node positions are updated based on relative positions and computed messages. Since the relative position vectors 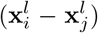 transform consistently under *E*(3) operations, this update preserves equivariance, meaning that if the positions are transformed (rotated or translated), the updated positions will transform in the same manner.

### Node Feature Update

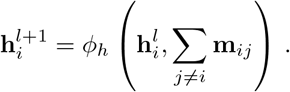

Node features are updated through a function *ϕ*_*h*_ that aggregates the messages from neighboring nodes. Since the feature updates depend only on invariant quantities like the aggregated messages, this ensures that the node feature transformations respect the symmetries of the input.

## F Input Representation

One of the first model design choices involves determining the resolution level at which we model the data. For peptides, which are a key focus of our work, there are two primary resolution levels: atom-level and residue-level modeling.

**Atom-Level Modeling** involves representing all heavy atoms (i.e., all atoms except hydrogen) as nodes in the peptide graph. This approach results in a higher number of nodes and increased computational cost. However, it captures more detailed structural information.

**Residue-Level Modeling**, alternatively, represents peptides at the level of amino acids, often referred to as residue-level representation. This method uses fewer nodes, reducing computational expense, but only provides one position per amino acid. For proteins, it has been demonstrated that the full structure can be predicted using only the locations of the C-alpha atoms. However, for our diffusion model, this approach has the disadvantage of providing less detailed input, as it only includes the C-alpha atom locations. On the other hand, it simplifies the peptide graphs, making the graph learning problem more tractable. Given that our work involves peptides conditioned on proteins, residue-level representation becomes especially important. Thus, we focus exclusively on residue-level representation in this work.

Beyond the basic 3D location of the C-alpha atoms, there are additional ways to enrich the input representation:

1. **Directional Information of Side Chains**: Incorporating the orientation of side chains can provide more detailed structural context. Utilizing an architecture that stores directional information along an additional axis in the hidden representations can be highly beneficial. For instance, architectures like Ponita (Bekkers et al. 2023) are designed to integrate such directional information, which could enhance the model’s performance by capturing more nuanced structural details.
2. **Rotation Matrix of the Backbone Frames**: Another alternative is to additionally input a representation of the rotation matrix of the backbone frames. This approach, used by AlphaFold 2 (Jumper et al. 2021), involves encoding the spatial orientation of the peptide backbone as a quaternion, offering additional structural context. A quaternion is a four-dimensional extension of complex numbers used to represent rotations in three-dimensional space. It is represented as *q* = *w* + *xi* + *yj* + *zk*, where *w, x, y*, and *z* are real numbers, and *i, j*, and *k* are the fundamental quaternion units. This representation avoids issues like gimbal lock that can occur with other rotation representations (Kuipers 1999). Integrating this information can significantly improve the model’s ability to predict and learn peptide structures by providing a richer representation of the peptide’s spatial configuration.

These alternatives to basic residue-level modeling offer the potential for more detailed and accurate predictions, making them valuable considerations for improving peptide modeling.

## G Positional Encoding

Positional encoding is used to incorporate information about the relative locations of each amino acid in the peptide chain. This helps the model transform peptide nodes, initially initialized from an isotropic Gaussian distribution, into the correct chain-like structure.

One common method for positional encoding is sinusoidal encoding (Vaswani et al. 2017). For a position *p* and dimension *d*, the sinusoidal positional encoding **PE**(*p, d*) is defined as:

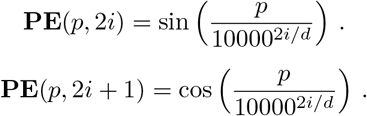

where *p* is the position in the sequence, *i* is the dimension index, and *d* is the dimensionality of the positional encoding. This encoding provides each position with a unique representation based on sine and cosine functions of varying frequencies.

In our model, positional encoding is added directly as extra feature dimensions to the peptide node embeddings.

Alternatively, positional encodings can be learned during training rather than using fixed sinusoidal functions. Learnable positional encodings offer the flexibility to adapt and optimize the encoding representation based on the specific learning task, potentially improving the model’s performance.

Overall, integrating positional encoding—whether fixed or learnable—ensures that the model can correctly interpret and utilize the spatial arrangement of amino acids within the peptide chain.

## H Noise Schedule and Sampling Path issue

### Noise Schedule

We employ a pre-defined polynomial noise schedule as introduced by Hoogeboom et al. (**?**). The noise schedule is defined as:

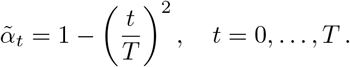

Different polynomial orders, such as linear, can be used in defining the noise schedule, but we have found that using a polynomial of order 2 is optimal for our model. Alternatively, the noise schedule can also be learned during training, allowing the model to adapt the schedule based on the specific requirements of the dataset and learning task.

Following Nichol & Dhariwal (Nichol et al. 2021) and Hoogeboom et al. (Hoogeboom et al. 2022), the values of

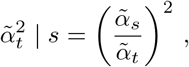

are clipped between 0.001 and 1 for numerical stability near *t* = *T*, and 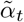 is recomputed as:

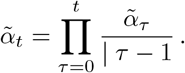

A small offset *ϵ* = 10^*−*5^ is used to avoid numerical problems at *t* = 0, defining the final noise schedule:

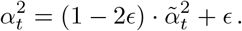

As mentioned in Section 3.1, 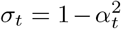 defines a variance-preserving noise schedule. However, this variance preservation only holds if the dataset variance is of the same scale as that of the starting distribution *p*(*z*_*T*_), which is not the case for our dataset. While it is possible to downscale the data or upscale the noise distribution, we have found that this makes the sampling path too hard to learn with the limited amount of data that we have.

### Sampling Path Issue

The mismatch in variance between the dataset and the starting distribution *p*(*z*_*T*_) leads to several issues. Given the scale of the data, the sampling path becomes very narrow. Additionally, the starting distribution has a much smaller variance than the data distribution, causing individual nodes to consistently expand from the origin outward to their target positions. This expansion introduces a bias into the model. As a result, if a node moves too far, it exits the learned sampling path and diverges from its target position. Our results demonstrate that it is possible to train a model that consistently produces good structures. However, due to this issue, achieving such results requires extreme precision in model construction.

## I COM for protein pocket and sampling initialization

Although the center of mass (COM) handling for the peptide is guided by the translational invariance argument, the treatment of the protein pocket still requires careful consideration.

In our model, the protein pocket is moved in accordance with the peptide’s center of mass, ensuring that their relative positions are maintained. This approach ensures that the entire joint p-MHC complex remains translationally invariant.

An alternative approach would be to center the MHC pocket at the origin and keep it fixed during the noising and generation processes. This might seem intuitive because it simplifies the learning problem, as the model does not need to learn to accommodate varying MHC positions for the same p-MHC. However, this method has the drawback of losing the correct relative position between the peptide and the protein pocket. While the relative position could be recovered using known anchoring points, this introduces additional complexity and effort.

Additionally, the initialization of the peptide needs to be considered in the context of COM handling. Given the subspace trick (related to peptide COM handling and translational invariance), we initialize the peptide at the origin using our starting noise distribution. We also center the protein pocket at the origin. This configuration places the peptide directly in the middle of the protein pocket, which might not represent the correct relative position but is the best we can achieve. The model can correct the relative position during sampling, but this does make the sampling start an unstable phase, where the model can fail to start following the noise path correctly.

## J Additional Information

### J.1 Code availability

The full code used for training, evaluation, and generation is publicly available at: https://github.com/DavidFruehbuss/MHC-Diff.

### J.2 Data availability

Both the 8K and 100K datasets, including clustering and data splits, will be made publicly accessible via Zenodo with a DOI upon acceptance of the manuscript for publication.

### J.3 Hardware

Experiments were conducted on NVIDIA A100 GPUs. A total of approximately 800 GPU hours were used for training, validation, and testing of the models.

### J.4 Author contributions

- **David Frühbuß**: Designed and developed the method, ran all experiments, wrote and edited the manuscript.
- **Coos Baakman**: Contributed to dataset construction and preprocessing code development.
- **Siem Teusink**: Provided prior project work; contributed to project and model design, including positional encoding development.
- **Erik Bekkers**: Supervised model design and project development; assisted in manuscript review and editing.
- **Stefanie Jegelka**: Supervised model design and project development; assisted in manuscript review and editing.
- **Li C. Xue**: Supervised project design and development; assisted in manuscript review and editing.

All authors reviewed and approved the final manuscript.

## Notes

https://github.com/DavidFruehbuss/MHC-Diff

### Competing Interest Statement

The authors have declared no competing interest.

